# Characterization of a novel zebrafish model of *MTMR5*-associated CMT4B3

**DOI:** 10.1101/2024.04.18.590157

**Authors:** Jordan Lindzon, Maia List, Salma Geissah, Mo Zhao, James J. Dowling

**Author notes:** These authors contributed equally to this work (Jordan Lindzon, Maia List).

## Abstract

Biallelic loss of expression/function variants in *MTMR5* cause the inherited peripheral neuropathy Charcot-Marie-Tooth (CMT) Type 4B3. There is an incomplete understanding of the disease pathomechanism(s) underlying CMT4B3, and despite its severe clinical presentation, currently no disease modifying therapies. A key barrier to the study of CMT4B3 is the lack of pre-clinical models that recapitulate the clinical and pathologic features of the disease. To address this barrier, we generated a zebrafish CRISPR/Cas9 mutant line with a full gene deletion of *mtmr5.* Resulting homozygous deletion zebrafish are born at normal Mendelian ratios and have preserved motor function. However, starting by 14 day-post-fertilization, mutant zebrafish develop obvious morphometric changes in head size and brain volume. These changes are accompanied at the pathological level by abnormal axon outgrowths and by the presence of dysmyelination, changes reminiscent of the nerve pathology in human CMT4B3. Overall, our *mtmr5* zebrafish mirror genetic, clinical, and pathologic features of human CMT4B3. As such, it represents a first pre-clinical model to phenocopy the disease, and an ideal tool for future studies on disease pathomechanism(s) and therapy development.

**Summary Statement:** We created a novel zebrafish *mtmr5/sbf1* mutant model of Charcot-Marie-Tooth Type 4B3 that recapitulates key features of the human disorder and provides the first *in vivo* model for therapy development.

## Introduction

Charcot-Marie-Tooth Disease Type 4B (CMT4B) is a rare, recessive subtype of inherited peripheral neuropathy (Previtali et al., 2007). Onset is in infancy or early childhood, and affected individuals experience progressive lower limb weakness and discoordination that ultimately leads to wheelchair dependence. There are a few therapeutic candidates from pre-clinical studies, and currently no treatments for CMT4B.

There are three genetic subtypes of CMT4B: CMT4B1 (due to biallelic mutations in *MTMR2*) (Bolino et al., 2004), CMT4B2 (due to biallelic mutations in *SBF2*/*MTMR13*) (Tersar et al., 2007), and CMT4B3 (due to biallelic mutations in *SBF1/MTMR5*) (Nakhro et al., 2013). All three subtypes share elements of peripheral nerve pathology (including characteristic myelin outfoldings), and all three are assumed to have common pathomechanisms, as the causative genes encode members of the myotubularin-related family of phosphoinositide phosphatases (MTMRs). Phosphatase active MTMRs (such as MTMR2) dephosphorylate PI3P and PI(3,5)P2, and are involved in the regulation of vesicle trafficking through the endolysosomal compartment (Bolis et al., 2007). Phosphatase inactive MTMRs are primarily thought to regulate the localization and/or activity of phosphatase active MTMRs. Both MTMR5 and MTMR13 directly interact with MTMR2 through their coiled-coil domains to form heterodimers, and both are proposed to function as regulators of MTMR2 activity (Kim et al., 2003, Robinson and Dixon, 2005).

CMT4B3 is unique as compared to CMT4B1/2 because patients experience signs and symptoms unrelated to peripheral demyelination. These unique clinical features include axonal peripheral neuropathy (Gang et al., 2020), structural brain changes (fork and bracket syndrome) (Romani et al., 2016), intellectual disability (Berti et al., 2021), and alterations in skeletal muscle. The expanded phenotype of CMT4B3 suggests that MTMR5 has functions beyond those specifically related to MTMR2, and also that therapies targeted at CMT4B1/2 may not fully address critical aspects of the CMT4B3 clinical syndrome.

To date, no pre-clinical models of CMT4B3 recapitulate the key features of the human disease. *Mtmr5* knockout mice have been generated; they have defective spermatogenesis (Firestein et al., 2002, Mammel et al., 2022), and peripheral axon demyelination without overt axon defects or brain involvement (Mammel et al., 2022). Zebrafish serve as an excellent alternative model organism for studying CMT4B3. They offer the advantage of rapid development, large numbers of offspring, easily detected and quantifiable motor phenotypes, and facile genetic manipulation (Santoriello and Zon, 2012). Zebrafish have previously been shown to accurately model other subtypes of CMT, particularly ones with axonal involvement (such as CMT2A) (Reddy et al., 2013), and to be useful for studying the function of MTMRs (Dowling et al., 2010) and myelination (Preston and Macklin, 2014). More importantly, zebrafish are an ideal organism for *in vivo*, whole animal drug screening (Karuppasamy et al., 2024), and are unique among tractable vertebrate model systems for this ability.

Similar to humans, zebrafish possesses one copy of the MTMR5 gene. MTMR5 proteins are highly conserved from zebrafish (NP_001038623.1) to human (NP_001352748.1) (72.13% identity), and both contain six functional domains (Fig. 1A): a tripartite DENN domain that regulates membrane trafficking by mediating Rab GTPase activity (Marat et al., 2011, Zhang et al., 2012), a SBF2 domain binding to SET-containing proteins that regulate chromatin structure (Cui et al., 1998, Dillon et al., 2005), two phosphoinositides or membrane binding domains consisting of a GRAM domain and a pleckstrin homology (PH) domain, a pseudo/inactive myotubularin phosphatase domain, and a coiled-coil domain that forms heterodimers with MTMR2 (Kim et al., 2003). Of note, the presence of N-terminal DENN domains is a unique feature of MTMR5 and MTMR13 in the myotubularin family (Laporte et al., 2003). Whilst missense CMT4B3 patient variants (Mégarbané et al., 2010, Nakhro et al., 2013, Alazami et al., 2014, Manole et al., 2016, Romani et al., 2016, Flusser et al., 2018, Gang et al., 2020, Berti et al., 2021) are clustered in the DENN domains and the SBF2 domain (Fig. 1A), suggesting their importance in normal function of MTMR5, pathogenic variants are reported throughout the gene (Landrum et al., 2020), and are associated with loss of expression and/or function (see Supplementary Table 1 for detailed information on the pathogenic variants).

**Figure 1.**
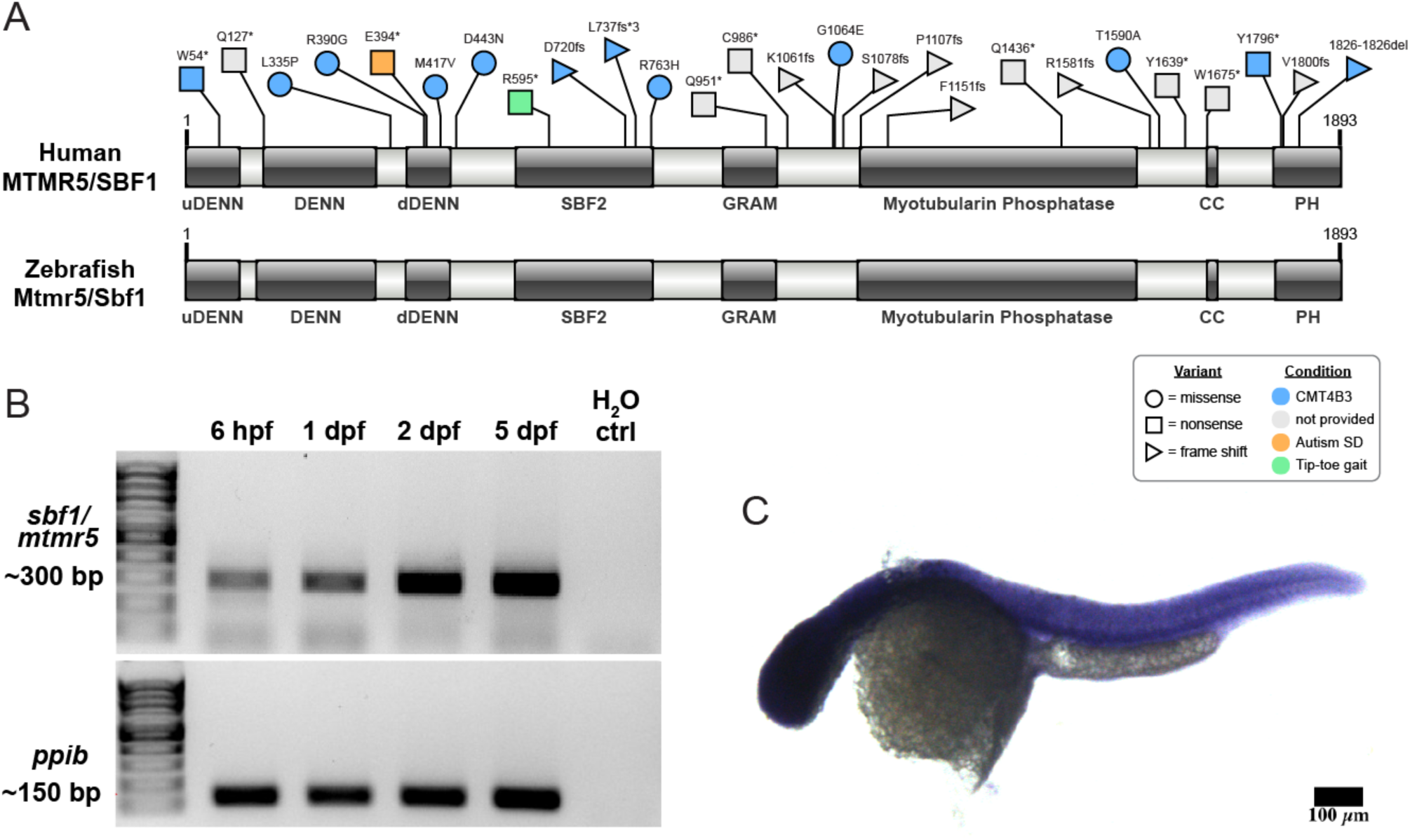
MTMR5 domain topology, disease variants, and spatiotemporal expression pattern in zebrafish. (**A**) MTMR5 contains six functional domains: an N-terminal tripartite Differentially Expressed in Neoplastic versus Normal (or DENN) domain [upstream DENN (a.a. 1-86), DENN (a.a. 129-311), and downstream DENN (a.a. 364-433)] often found in proteins involved in Rab-mediated membrane trafficking; a SBF2 (SET binding factor) domain (a.a. 543-765) that interacts with SET-domain-containing proteins; a GRAM domain (a.a. 883-969) that is a lipid/intracellular protein binding domain; a catalytically inactive myotubularin-phosphatase domain (a.a. 1108-1533); a coiled-coil domain (a.a. 1651-1665) that mediates binding with MTMR2; and a C-terminal PH (pleckstrin homology) domain (a.a. 1763-1868) that mediates binding with phosphoinositides. Pathogenic missense variants in MTMR5 mostly cluster at the dDENN and SBF2 domains, whereas hypomorphic variants (nonsense, frameshift, etc) are found throughout the gene (details see Supplementary Table 1). Of note, the domain topology of the MTMR5 protein is highly conserved between human and zebrafish. Autism SD: autism spectrum disorder. (**B**) RT-PCR followed by an agarose gel (1%) electrophoresis shows the presence of *mtmr5* in total zebrafish cDNA at 6 hour-post-fertilization (hpf), 1 day-post-fertilization (dpf), 2 dpf, and 5 dpf, but not in the water (H_2_O) control. Note: *ppib* is a housekeeping control. (**C**) Whole-mount in situ hybridization using DIG-conjugated RNA probes in 1dpf embryos shows that *mtmr5* is ubiquitously expressed. Scale bar: 100 μm. Domain topology of MTMR5 proteins was mapped using SMART (Letunic et al., 2021) and DeepCoil (Ludwiczak et al., 2019). Domain illustrations were generated using IBS 2.0 (Xie et al., 2022).

In this study, we generated and characterized a novel zebrafish model of MTMR5-associated CMT4B3. We established a zebrafish *mtmr5* knockout (KO) line via CRISPR/Cas9-mediated full gene deletion, and confirmed the absence of *mtmr5* mRNA transcripts in the resulting homozygous mutants. Phenotypic characterization revealed that *mtmr5*-KO zebrafish recapitulate many key features of CMT4B3, including axonal defects, dysmyelination, and microcephaly. In total, our findings establish *mtmr5* mutant zebrafish as an ideal model for studying MTMR5-associated CMT4B3 pathomechanisms and for identifying potential therapies.

## Results

### MTMR5 is ubiquitously expressed in zebrafish

To examine the spatiotemporal expression pattern of *mtmr5* in zebrafish, we performed RT-PCR and whole mount *in situ* hybridization. We detected *mtmr5* mRNA as early as 6 hours post fertilization (hpf) in zebrafish (Fig. 1B) and across all tissues at 1 day post fertilization (dpf) (Fig. 1C), confirming early and ubiquitous expression similar to mammalian models.

### Generation of a zebrafish *mtmr5* knockout model via CRISPR/Cas9 genome editing

In order to study MTMR5 function, we generated a zebrafish line with a full gene deletion of *mtmr5,* i.e. a ∼86 kb region (spanning from 40 bp downstream to translation start site, to 3 bp downstream to stop codon) was removed from the genome (Fig. 2A-C). Briefly, we designed and tested two highly efficient guide RNAs (gRNAs) against the zebrafish *mtmr5* gene, one cutting near the translation start codon (ATG), and the other near the stop codon (TAG). gRNAs were co-injected with Cas9 mRNA into 1-cell stage wild-type embryos, and injected embryos were raised to adulthood. Primers flanking the *mtmr5* gene were used to screen adult founders (F0) via PCR for carrying a ∼500 base pair (bp) band. The genotype was further confirmed by Sanger sequencing.

**Figure 2.**
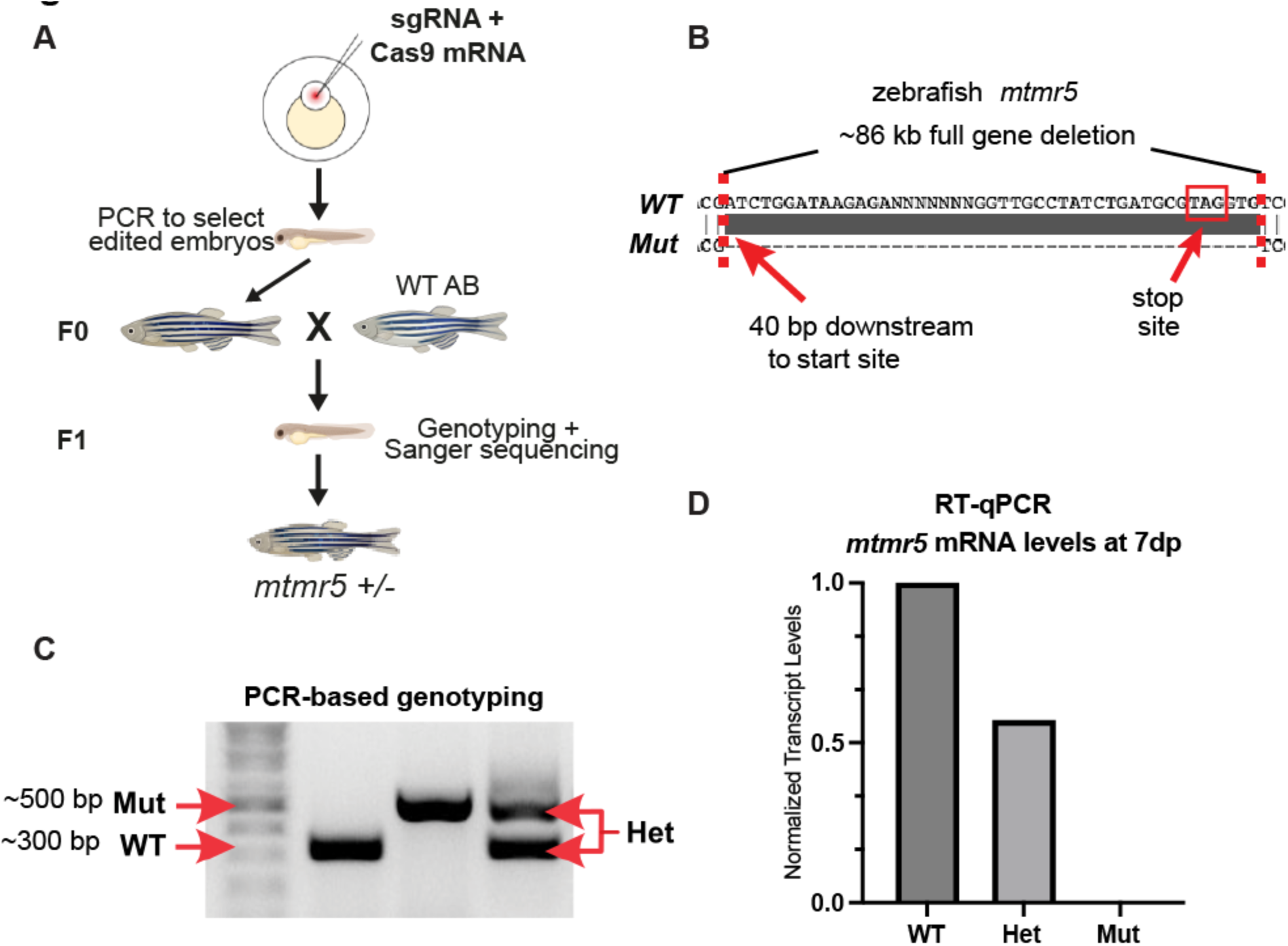
Generation of *mtmr5* knockout zebrafish. (**A**) Schematic displaying the workflow used to generate the *mtmr5* mutants. Highly efficient guide RNAs targeting zebrafish the 5’ and 3’ UTRs of *mtmr5* were co-injected with Cas9 mRNA into one-cell-stage WT (AB strain) embryos. Injected embryos were raised to adulthood, outcrossed to WT, and founders (F0) were identified by screening resulting embryos for gene deletion using PCR-based genotyping. Founders were outcrossed to WT, and embryos with the desired deletions were raised to adulthood (F1 generation). The genotype of F1 was then confirmed by PCR and Sanger sequencing. (**B**) Sanger sequencing on genomic DNAs from F1 adults revealed an 86 kilobase (kb) deletion starting 40 base pairs (bp) downstream of the MTMR5 translation start site to 3 base pairs downstream of the translational stop site. (**C**) PCR-mediated genotyping using genomic DNAs were resolved by 1% agarose electrophoresis after PCR. Three primers were used: a common forward primer with a mutant-specific reverse primer that flanks the entire *mtmr5* gene (upper band), and with a wild type-specific primer that binds to an internal region of the gene (lower band). Mutant amplicon was observed when the deletion was present (mutants and heterozygotes) and Wild type amplicon were observed when there was no deletion (WT and heterozygotes). (**D**) qRT-PCR analysis shows significant reduction (∼50%) of *mtmr5* expression in the heterozygotes and no expression in the mutants at 7 dpf.

To demonstrate the molecular consequence of this mutation, we performed RT-qPCR, and found an absence of *mtmr5* mRNA in the zebrafish homozygous for the deletion mutation [i.e. *mtmr5* mutants or *mtmr5* knockouts (KOs)] (Fig. 2D). We thus successfully generated an *mtmr5* KO zebrafish line via CRISPR/Cas9-mediated full gene deletion.

### Loss of *mtmr5* is associated with early-onset microcephaly, and overall size reduction upon adulthood, but no changes in survival or motor function

*mtmr5* homozygous mutant zebrafish are viable and can survive to adulthood. To characterize *mtmr5* loss-of-function effects on neuromuscular function, we performed swim assays on 3dpf, 6dpf, and 14dpf zebrafish using ZebraBox (Viewpoint). ZebraBox is a semi-automatic platform that can be used to track the movement of individual embryo/larva in 96-well plates. We first compared distance travelled by wild-type siblings (WT) versus KOs generated from heterozygous in-crosses, and observed no difference (Fig. 3A-C). In order to remove the contribution of maternally deposited *mtmr5*, we further characterized offspring from homozygous females crossed to heterozygous males, and again observed no difference in swim performance (Fig. 3D-E). In all, the loss of *mtmr5* does not affect survival or cause significant movement defects, similar to observations from *Mtmr5* -/- mice (Mammel et al., 2022).

**Figure 3.**
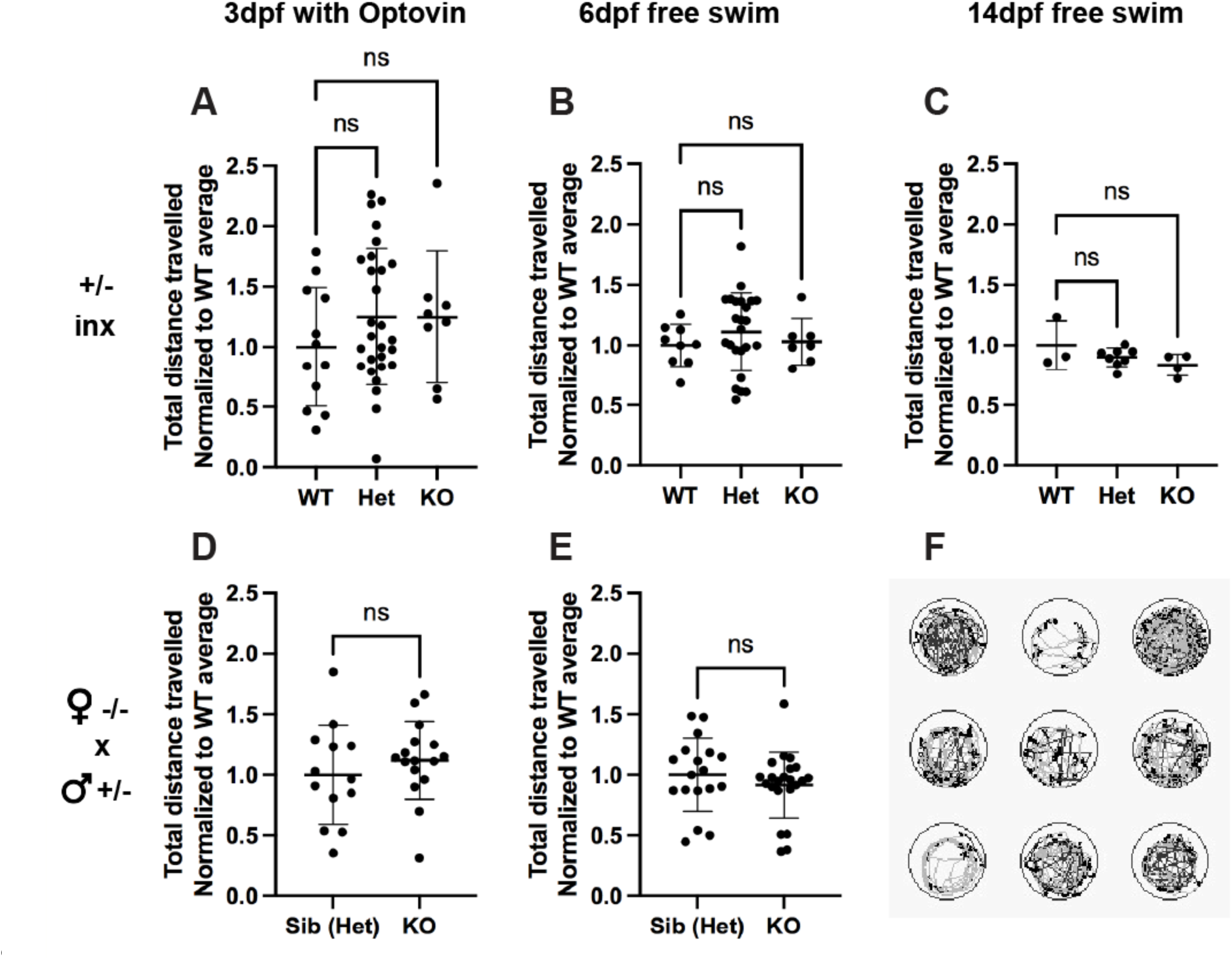
*Mtmr5* mutants have normal swim behavior. (**A**) Embryos generated from a heterozygous in-cross were subjected to Optovin treatment at 3 days post fertilization (dpf) followed by the quantification of the total distance travelled by each embryo via Zebrabox. There was no apparent difference in swim performance between wild type (WT), heterozygous (Het), and knockouts (KO). (**B, C**) Embryos resulting from a heterozygous in-cross were subjected to a free swim assay (no Optovin treatment) at 6dpf (B) and 14dpf (C). There were no observed differences in swim performance between the three genotypes at both developmental stages. (**D, E**) Embryos resulting from a homozygous knockout female crossed with a heterozygous male were subjected to swim assays at 3dpf (with Optovin treatment) (D) and at 6dpf (free swimming assay) (E), and again, at both developmental time points, there were no differences in swimming performance between the genotypes. (**F**) A representative image of the swim trace tracking used by the Zebrabox program to quantify the total distance travelled for each embryo per well. All data were normalized to the average of controls (WT or sibling heterozygous). All statistical analyses include at least three independent experiments. Each dot on the graphs represents one zebrafish embryo, at least n=4 embryos were used per group per experiment. Data are mean +/-s.e.m. One-way ANOVA was done for all heterozygous in-cross experiments and an Unpaired two-tailed Student’s t-test was used for the HomFxHetM experiments. Ns = not significant, P> 0.05.

We then characterized the gross morphology of *mtmr5*-KO zebrafish by measuring brain/body size. At 14 dpf (but not before), we noted in KOs significant reduction in brain height, as well as reduction in the ratio of brain to body height (Fig. 4A-F), indicating the presence of a small brain phenotype (i.e. microcephaly). In addition, at 2 months of age, *mtmr5*-KO showed significant reduction in both body length and brain height (now with normal brain/body ratio) (Fig. 4G-L). The reduced overall body size is consistent with smaller body size reported in *Mtmr5* -/-adult mice (Mammel et al., 2022).

**Figure 4.**
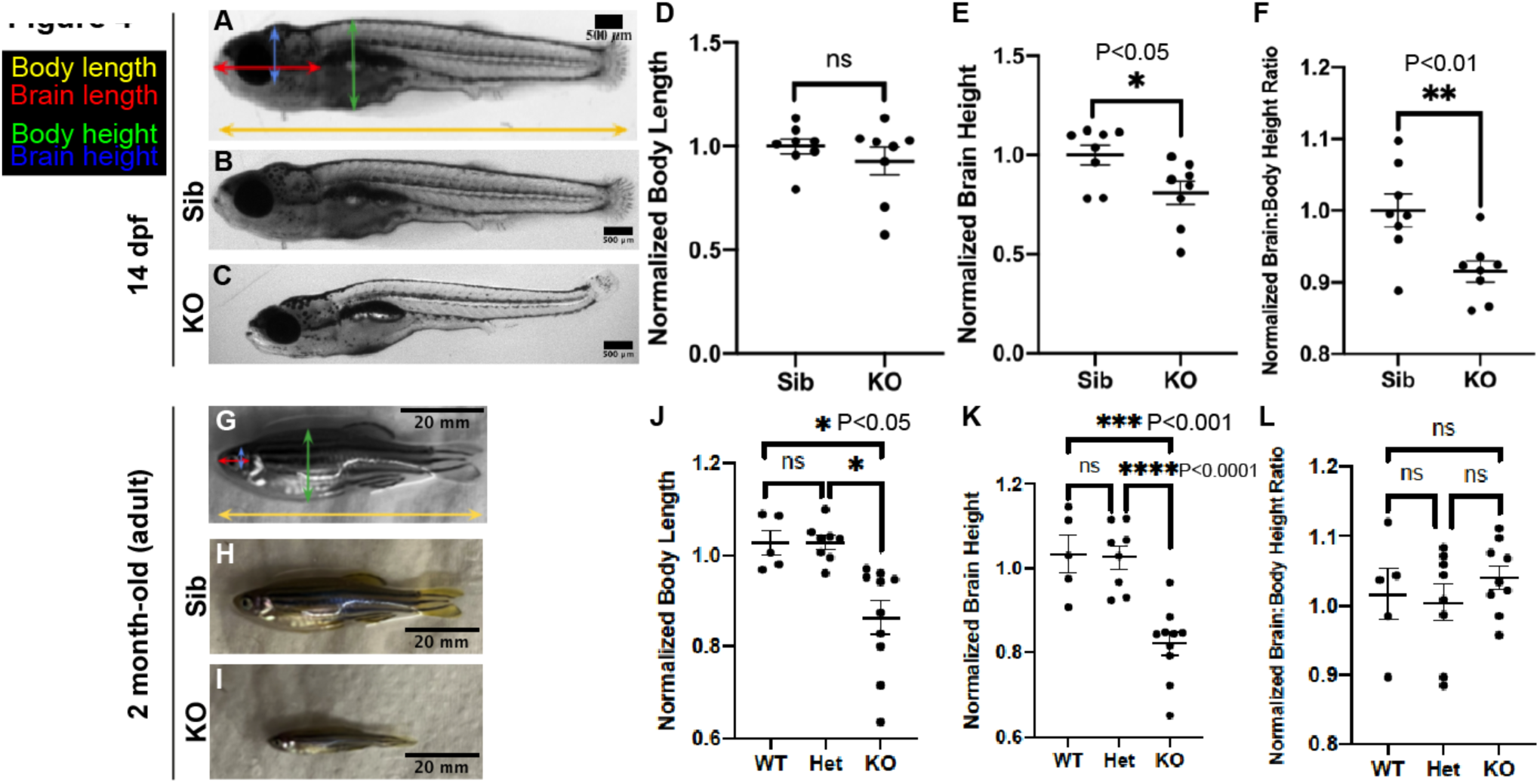
Loss of MTMR5 results in a gross morphology phenotype, starting with microcephaly 14 dpf and leading to an overall body size reduction. (**A**) A representative image of a wild-type (WT) 14-day post-fertilization (dpf) zebrafish with lines delineating the measured gross anatomy. The yellow line spanning the entire zebrafish denotes the body length, and the red line spanning the lateral head region represents the brain length. The green line spanning from the beginning of the pectoral fin to the dorsal end of the fish is the body height. Brain height is delineated by the blue line starting at the ventral end of the eye to the dorsal end of the brain. (**B**) A representative image of a WT sibling (sib = WT) at 14dpf and (**C**) a representative image of a knockout (KO) at 14dpf where a decrease in brain size is observed. Lateral images were taken of individual zebrafish, and measurements were done via the Line Tool on Fiji ImageJ, and then normalized to the average sibling measurement. (**D**) There was no observed difference in body length between WT and KO at 14dpf. (**E**) Brain height was significantly reduced in the KOs compared to WT and (**F**) the brain-to-body height ratio was similarly reduced in the KOs. (**G**) A representative image of a WT 2-month-old (adult) zebrafish with lines delineating the measured gross anatomy. Lines drawn on the zebrafish describe the same parameters measured in Figure A. (**H, I**) Representative images of WT and KO adult zebrafish, where a decrease in KO total body size is observed. (**J-L**) Measurements were done similarly to 14 dpf. (**J**) Body length was found to be significantly decreased in the adult KO compared to both WT and heterozygous (Het) adult zebrafish. There were no observed differences between the adult WT and Het animals. (**K**) There was a significant reduction in brain height in adult KOs vs WT and Hets. (**L**) There was no observable difference in the brain-to-body height ratio between any of the genotypes at the adult stage. All statistical analyses include at least three independent experiments. Each dot on the graphs represents one zebrafish, at least n=5 zebrafish were used per group per experiment. Data are mean +/-s.e.m. Unpaired two-tailed Student’s t-test was used for the 14 dpf measurements, and One-way ANOVA was done for all 2-month-old measurements. *P<0.05; **P<0.01; ****P<0.0001; ns, not significant. Scale bars: 500 μm (14dpf images) or 20 mm (adult images).

We further examined the microcephaly phenotype to determine if it was due to changes in aspects of nerve morphology or instead excessive cell death. To study apoptosis in the nervous system, we performed acridine orange (AO) staining on live zebrafish at 7 dpf (Fig. 5A) and 14 dpf (Fig. 5C-D), but did not observe in the head (CNS, central nervous system) or in the trunk (PNS, peripheral nervous system) any accumulation or overall increase of AO positive cells. To visualize the ventricular system, we performed DAPI staining on fixed 7dpf zebrafish (Fig. 5B), and detected no differences between WT sibling and KO in the size of DAPI-free areas. In other words, KO caused no enlargement in brain ventricle size, which might be expended in there was brain atrophy due to brain cell loss. In total, these data demonstrate that the microcephaly phenotype in *mtmr5-*KO zebrafish is not caused by excessive apoptosis in the nervous system, indicating instead potential defects in early brain development.

**Figure 5.**
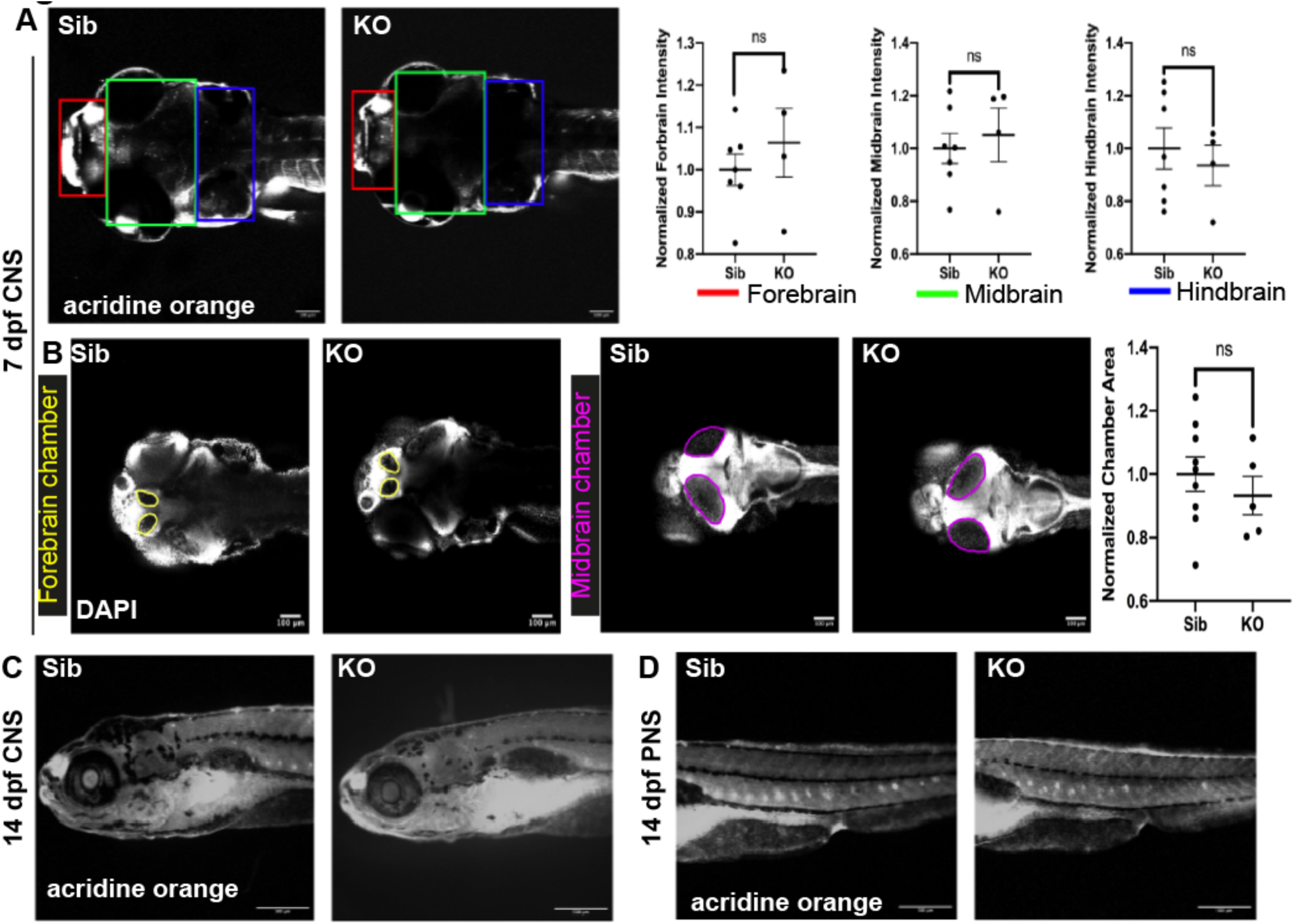
Loss of MTMR5 does not result in excessive cell death in the nervous system and does not cause changes in brain ventricle size. (**A**) Representative average Z-projections depicting dorsally orientated 7 dpf zebrafish embryos stained with acridine orange (AO), a live stain used to visualize and quantify apoptotic cells. The red box (most leftward) denotes the forebrain, the green box (middle) denotes the midbrain, and the blue box (most rightward) denotes the hindbrain. Using Fiji ImageJ, the intensity of AO staining in the forebrain, midbrain, and hindbrain was quantified. There was no statistically significant change in AO intensity in any of the brain regions between WT siblings (sib) and KO. (**B**) Representative images from fixed and DAPI-stained 7dpf zebrafish embryos, used to visualize and quantify brain ventricular chamber size. The forebrain is outlined in yellow (two left images) and the midbrain is outlined in magenta (two right images). There were no qualitative or quantitative differences in forebrain or midbrain ventricle size between sibs and KOs at 7dpf. (**C**) Representative live images of AO-stained anterior lateral regions of 14dpf zebrafish, encompassing the central nervous system, show no visual differences between sibs and KOs. (**D**) Representative live images of AO-stained posterior-lateral regions of 14dpf zebrafish, encompassing the peripheral nervous system, show no visual differences between sibs and KOs. All statistical analyses include at least three independent experiments. Each dot on the graphs represents one zebrafish, at least n=3 zebrafish were used per group per experiment. Data are mean +/-s.d. Unpaired two-tailed Student’s t-test was used: ns, not significant. Scale bars: 100 μm

### Loss of *mtmr5* induced axon defects and dysmyelination in the developing peripheral nervous system

CMT4B3 patients showed both axon and myelination defects in the peripheral nervous system (Alazami et al., 2014, Romani et al., 2016, Flusser et al., 2018). In zebrafish, while all major components of the brain are present by 5 dpf, primary motor neurons continue to develop until 30 dpf and the lateral line (composed of peripheral sensory neurons) is not fully developed until 2 months (Cunliffe, 2003). To examine axon patterning in *mtmr5*-KO embryos, we performed wholemount immunofluorescence on 7dpf zebrafish larvae using an antibody against acetylated tubulin, which labels mature axons. Interestingly, confocal images of KOs showed disorganized and less defined axon morphology (Fig. 6A-B). To quantify axon branching, we utilized the Skeleton plugin in Fiji ImageJ (Fig. 6A’-B’), which measures parameters such as the number of axon branches, the number of end-points (where branches end), and the length of axon branches. We detected in the KOs a significant upregulation of the total number of branches (Fig. 6C) and the total number of end-points (Fig. 6D), while the average branch length was significantly reduced (Fig. 6E). These data support that *mtmr5* KO may disrupt axon path finding and/or growth during neural development.

**Figure 6.**
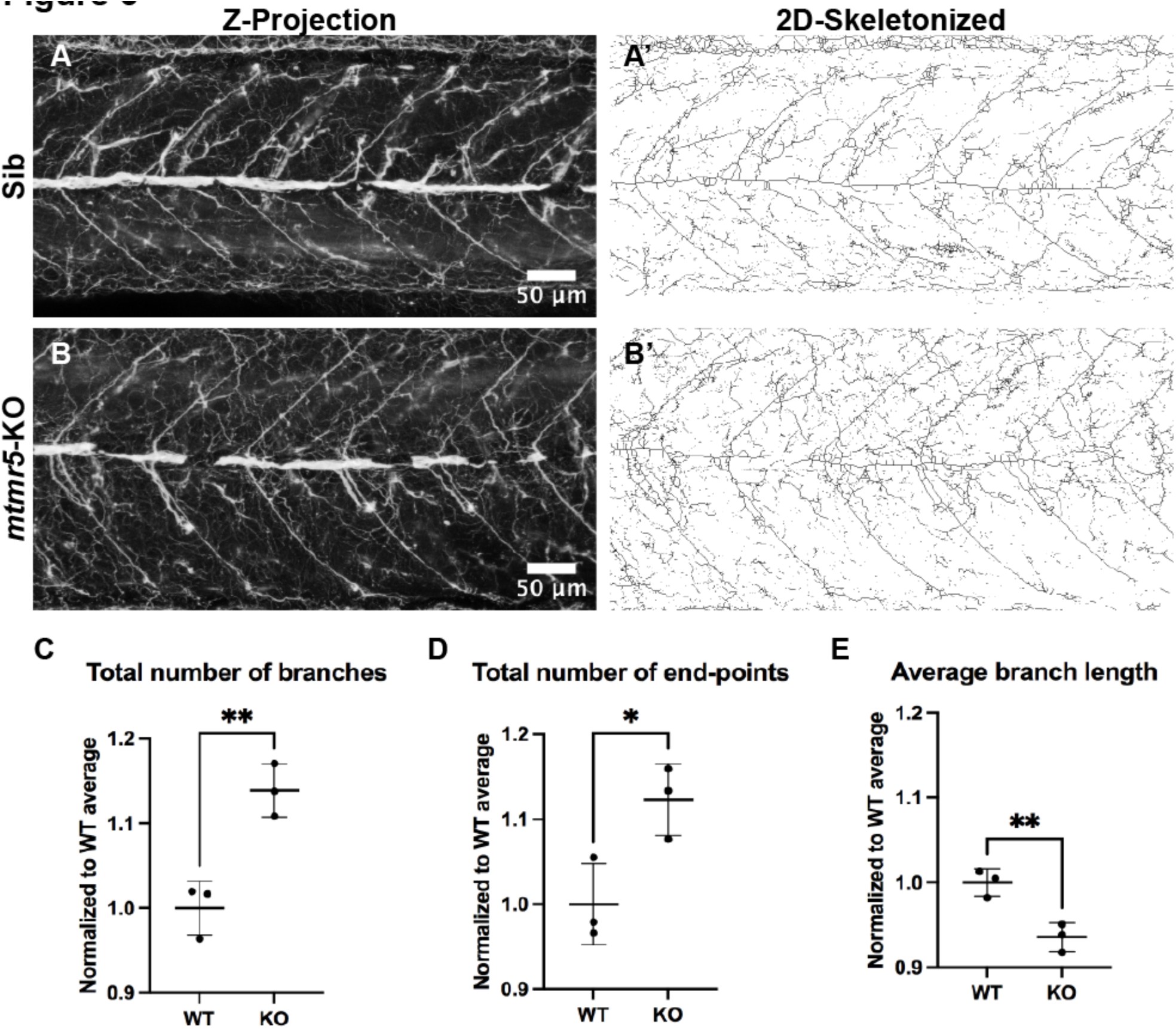
MTMR5 loss leads to less defined axonal organization with increased total number of axon branches and decreased branch length. (**A-B**) 7 dpf zebrafish embryos were fixed and probed with an acetylated tubulin antibody to label mature axons. (**A, B**) Representative images of Z-projections (standard deviation via Fiji ImageJ) of acetylated tubulin staining. Qualitatively, axons appear less defined in the knockout (KO) compared to the sibling. (**A’, B’**) 2D skeletonized images generated using Binary function in Fiji ImageJ to allow for quantification of axonal organizational parameters. All quantification was normalized to the wild-type (WT) average of the respective parameter. The number of total axonal branches (**C**) and axonal endpoints (**D**) were significantly increased in the KO compared to the WT. (**E**) The average axonal branch length was significantly decreased in the KO compared to the WT. All statistical analyses include at least three independent experiments. Each dot on the graphs represents one zebrafish, n=3 zebrafish were used per group per experiment. Data are mean +/-s.d. Unpaired two-tailed Student’s t-test was used. *P<0.05; **P<0.01, ns, not significant. Scale bars: 20 μm.

To examine PNS myelination in the KO zebrafish, we imaged transverse sections of WT versus KO posterior lateral line under transmission electron microscopy. In 14dpf WT siblings, most axons were in close proximity to the myelin sheaths (Fig. 7A-A’). In the KO, we did not observe lack of myelination, but did observe an increased number of axons showing plasma membrane detachment from the myelin sheaths (Fig. 7B-B’), consistent with a dysmyelination phenotype.

**Figure 7.**
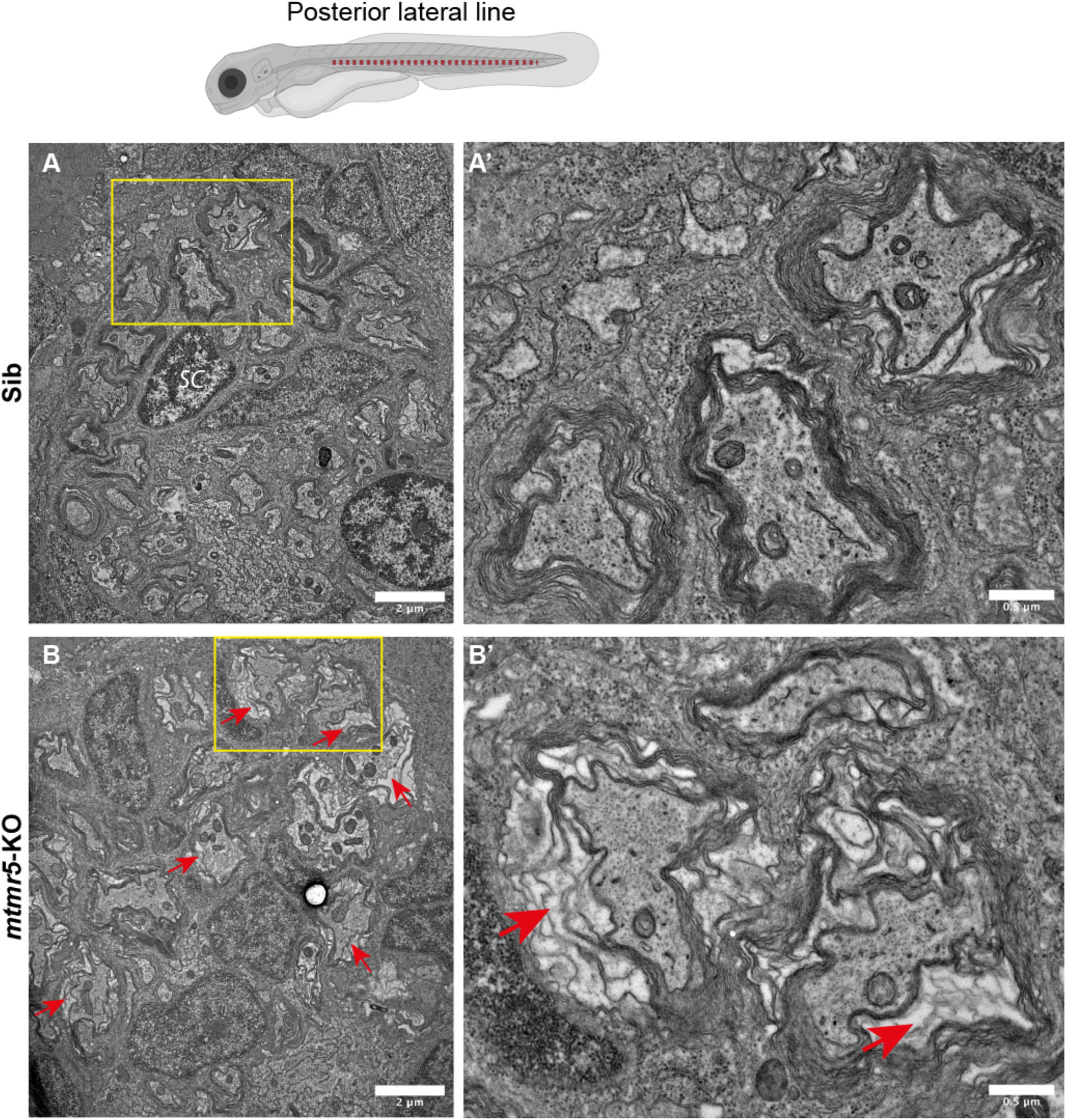
Loss of MTMR5 results in dysregulated axon myelination. Transmission electron microscopy was used to visualize the posterior lateral line (PLL) in 14 dpf zebrafish. The PLL region is contextualized by the red dotted line on the zebrafish embryo schematic. **(A**) The axons in the 14 dpf WT siblings were in close contact with myelin sheaths. This tight and organized wrapping can be observed in more detail in the zoomed-in inset (yellow box in A, full image in A’). (**B**) In the knockouts (KO) at 14dpf, the myelin is still present, yet there is an apparent white space between the axon and the myelin. This space is indicated by the red arrows. (**B’**) This is a zoomed-in version of the inset (yellow box) outlines in B, showing the detachment or loose myelin in the KOs. Scale bars: (A, B) 2 μm or (A’, B’) 0.5 μm.

## Discussion

CMT4B3 is an early-onset neuropathy that currently has no treatment. It is caused by recessive pathogenic variants in the *MTMR5* gene. In this study, we report the first *mtmr5* zebrafish knockout line generated by CRISPR/Cas9-mediated gene editing. Consistent with patient phenotype(s), loss of MTMR5 in zebrafish results in early-onset microcephaly without affecting overall survival or motor performance. Of note, brain or CNS involvement is a disease hallmark of CMT4B3 patients that has not been reported in other CMT4B subtypes or in the mouse models of MTMR5. Upon examining the molecular phenotypes in the nervous system, we observed disrupted axon patterning and dysmyelination in the PNS (peripheral nervous system) of *mtmr5-*KO zebrafish, similar to the polyneuropathy reported in patients (Mégarbané et al., 2010, Manole et al., 2016, Flusser et al., 2018). Our zebrafish model is thus a first *MTMR5* animal model that recapitulates multiple aspects of the clinical spectrum of CMT4B3 disease.

MTMR5 is a pseudophosphatase that binds to its active family member MTMR2 (Mammel et al., 2022). MTMR2 subcellular localization is either diffuse or cytoplasm-specific depending on cell type (Previtali et al., 2007), and its interaction with MTMR5 (Kim et al., 2003). MTMR5 thereby may function in the PNS (at least partially) by recruiting MTMR2 to its substrates and promoting its enzymatic activity (Kim et al., 2003). Notably, CMT4B1-causing *MTMR2* mutations drastically reduced its phosphatase activity (Berger et al., 2003), suggesting the importance of MTMR2 activity in ensuring normal PNS development/function.

MTMR5 may also have functions beyond those specifically related to MTMR2 and MTMR13. Each MTMR is associated with distinct patient neuropathies, and distinct nerve pathologies in animal models. *Mtmr2* -/- and *Mtmr13 -/-* mice showed classic CMT4B myelin outfolding in the PNS (Azzedine et al., 2003, Tersar et al., 2007, Robinson et al., 2008), whilst adult *Mtmr5 -/-* mice showed reduced number of myelinated axons likely due to disruption in axon radial sorting (Mammel et al., 2022). Our 14dpf *mtmr5* -/- zebrafish larvae show dysmyelination at the axon plasma membrane, in keeping consistent with both the *Mtmr5* KO mice and patient pathology. Furthermore, the expanded CNS phenotype of CMT4B3 patients and the microcephaly phenotype observed in our 14dpf *mtmr5-/-* zebrafish larvae support a novel role of MTMR5 in CNS development. Notably, a recent report identified MTMR5 association with other neurological disorders such as Alzheimer’s disease in aged population (Khamse et al., 2022), emphasizing a potential role in CNS maintenance and its degenerative processes.

At present, though, little is known about the importance of MTMR5 in neural development and brain homeostasis, as well as its subcellular localization/function in different neural cell types. Due to the absence of excessive cell death, these early neural phenotypes are likely a result of developmental delay rather than neurodegeneration. Future work is needed to unravel the role(s) of MTMR5 in brain development, and establish why its loss causes microcephaly. Moving forward, our *mtmr5* mutant zebrafish are ideal to explore the CNS function of MTMR5, particularly as the larvae manifest an obvious brain phenotype that mirrors the human disease and is not seen in the mouse KO model. Multi-omic platforms, such as spatial transcriptomics and proximity labeling with BioID, are potential approaches to unbiasedly identify disease pathways and interacting partners associated with MTMR5.

Lastly, we identified the microcephaly present at 14 dpf as the most feasible, and disease relevant phenotype for future therapy identification and development. Zebrafish serve as an excellent model for drug screens due to their high gene conservation with human and ease of drug administration (Karuppasamy et al., 2024). Candidate treatments to test include drugs previously shown to work in other CMT subtypes (Pisciotta et al., 2021, Stavrou et al., 2021) or other MTMR-related neuromuscular disorders, e.g. valproic acid (Volpatti et al., 2022), tamoxifen (Maani et al., 2018), and wortmannin (Sabha et al., 2016) in X-linked myotubular myopathy due to *MTM1* mutation. Candidates may also be identified from drug libraries targeting MTMR5-related pathways, for example endocytosis and autophagy (Chua et al., 2022).

## Materials & Methods

### Zebrafish maintenance and collection

Zebrafish (AB strain) were raised and maintained at 28.5°C at the Zebrafish Facility at the Hospital for Sick Children, Toronto, ON, Canada. Experiments were performed on zebrafish embryos and larvae (grown in blue water, 0.3 g/L Instant Ocean, 1 mg/L methylene blue, pH 7.0) from the one-cell stage up to 14 dpf. All zebrafish procedures were performed in strict accordance with the Animals for Research Act of Ontario and the Guidelines of the Canadian Council on Animal Care.

### RNA extraction, cDNA synthesis, and PCR

Total mRNA was isolated from 6 hpf to 5 dpf zebrafish homogenates using an RNeasy Mini Kit (Qiagen, 74106) and reverse-transcribed with iScript (Bio Rad, 1708891). Approximately 10-25 embryos were collected for each stage. PCR was performed using GoTaq Green Master mix (Fisher Scientific, PR-M7123) and a PCR cycler (Applied Biosystems). The zebrafish gene beta actin, *βact*, was used as the endogenous control. Primers used are as follows: *mtmr5* forward 5′-AAGCATCAGAACATCTGCCG-3′, reverse 5′-TGTTTTTGGCAGAGACAAGAAGT-3′; and *βact* forward 5′-CCATCCTGCGTCTGGATCTGGCTG-3′, reverse 5′-CGCCATACAGAGCAGAAGCCATG-3′; *ppib* forward 5’-ACCCAAAGTCACGGCTAAGG-3’, reverse 5’-CTGTGGTTTTAGGCACGGTC-3’. Protocol conditions for PCR were the following: denaturation at 95°C for 5 min; annealing at 58°C for 30 sec, extension at 72°C for 45 sec, followed by 40 cycles; and final extension at 72°C for 5 min.

### Wholemount *in situ* hybridization

DNA template was generated via pGEM T-easy cloning system (Promega, A1360), where the insert was generated by PCR from 2 dpf total cDNA synthesized using iScript (Invitrogen, 11755050). Primers were designed to target a ∼1 kb region at the 3’UTR of zebrafish *sbf1/mtmr5* mRNA transcript: forward 5’-CCTCATAGCCAATGGGGAGC-3’, and reverse 5′-GCTGGATATCGGAAGCGGAT-3′. Digoxigenin (DIG)-labeled *in-situ* probes were synthesized using DIG RNA Labeling Kits (Roche, 11277073910). RNA *in-situ* hybridization was carried out as previously described (Thisse and Thisse, 2007). Briefly, 1 dpf AB embryos were fixed in 4% paraformaldehyde (PFA) for 2 hours at room temperature, and then dehydrated in 100% methanol at -20°C until needed. Embryos were then permeabilized using Proteinase K (Thermo Scientific, EO0491) and incubated with DIG-labeled antisense RNA probes in hybridization solution. Hybridizations of the probe with the RNA were detected with an alkaline phosphatase-conjugated antibody (1:5000; anti-DIG-AP, Fab Fragments, Roche, 11093274910). Finally, stained embryos were cleared overnight in a gradient ethanol / PBST (0.1% Tween 20 in PBS) wash and imaged under a ZEISS AxioZoom stereomicroscope.

### CRISPR/Cas9 genome editing and full gene deletion

The program Chopchop (http://chopchop.cbu.uib.no/) (Montague et al., 2014) was used to design each of the guide RNAs (gRNAs) used in this project. Next, 50-100 one-cell-stage WT embryos were injected with the gRNA (150 pg per embryo) and Cas9 mRNA (100 pg per embryo) with a Picopump (World Precision Instruments). Genomic DNA was extracted using 50 μL of NaOH solution (50 mM) at 95°C for 20 min, and incubated at 4°C for 10 min, followed by neutralization using 50 μL of Tris-HCl (1M, pH 8.0). gRNA efficiency was determined using HRM analysis performed on a Roche Lightcycler 96. Once highly efficient gRNAs was identified at the desired genomic region, potential founders (F0) were outcrossed to WT AB zebrafish. In-cross progeny from the F3 and F4 generations were used for the characterization of the *mtmr5* mutant phenotype. To create full gene deletion mutants, the gRNAs used in this study were: gRNA (left) 5′-CGTTGTAGTCGGCTACGATC-3′ at exon 1, and gRNA (right) 5′-CCTATCTGATGCGTAGGTGT-3′ at exon 42 (last exon). Primers used for determining cutting efficiency via HRM were as follows: gRNA (left) forward 5′-TGGAAAGACAGTCATCCCCG-3′, reverse 5′-GCTGGCTGTGTCTCACTCTTA-3′; gRNA (right) forward 5′-CTCAAGACGACCAAAAGAGTGT-3′, reverse 5′-GCGTCACAGTTATGTTGTCTGC -3′. Primers used for determining full-gene deletion via PCR were as follows: common forward 5′-TATCTTCGTGCACGCGCTGTAA-3′, wildtype-specific reverse 5′-GCTGGCTGTGTCTCACTCTTA-3′ (328 base-pair product when paired with common forward); deletion-mutant-specific reverse 5′-GAAGGAAGTGTTTTGAAGCCGA-3′ (499 base-pair product when paired with common forward).

### Reverse transcription and quantitative PCR

Total RNA was isolated from 7 dpf *mtmr5* wildtype, heterozygous, and deletion mutant zebrafish larvae using the Monarch Total RNA Miniprep Kit (New England Biolabs (NEB, T2010S) and reverse-transcribed with High-Capacity cDNA Reverse Transcription Kit (Applied Biosystems, 4368814). Approximately 10-25 embryos were collected for each genotype. Quantitative PCR was performed using Luna Universal qPCR Master Mix (NEB, M3003L) and a Roche LightCycler 96 machine (Roche). The zebrafish βactin and GAPDH genes were used as housekeeping controls (βactin forward 5’-CGAGCTGTCTTCCCATCCA-3’, reverse 5’-TCACCAACGTAGCTGTCTTTCTG-3’; and GAPDH forward 5’-CGCTGGCATCTCCCTCAA-3’, reverse 5’-TCAGCAACACGATGGCTGTAG-3’). Gene-specific primers: *mtmr5* forward 5 ′ - CAGCGGATTCACACCAGTCT-3 ′, reverse 5’-CGGAAACGGTTGGAGACGTA-3’. Protocol conditions for quantitative PCR were the following: denaturation at 95°C for 60 sec; followed by 45 cycles of 95°C for 15 sec, 60°C for 30 sec.

### Gross morphology measurements

Genotyped zebrafish were treated with tricaine (1:25 in embryo water) and imaged under AxioZoom stereoscope (ZEISS). Brightfield images were then analyzed using Line Tool in Fiji ImageJ to measure brain length, brain height, body length, and body height. All measurements were normalized to the average of the sibling, and entered into Prism 9 (GraphPad). Student’s t-test was performed to analyze the statistical significance.

### Swim assay

To quantify muscle performance, 3-14 dpf zebrafish were individually transferred to a 96-well plate. 3 dpf zebrafish were incubated in an optovin analog 6b8 (10 µM in 200 µl embryo water, ChemBridge, 5707191) at 28.5°C for 5 min in the dark. Motor activity of the larvae was recorded and analyzed using ZebraBox (Viewpoint, France) as previously described (Zhao et al., 2019) with 30 sec light on, 1 min light off, 30 sec light on, 1 min light off and 30 sec light on. Without optovin incubation, 6 dpf or 14 dpf zebrafish were tracked in the 96-well plate in the ZebraBox (dark) for free swim for 2 hours at room temperature. Three independent experiments were conducted. Total distance traveled (mm) was plotted and analyzed using Prism 9 (GraphPad). For each group, S.E.M. was calculated, and one-way ANOVA was performed to test statistical significance.

### Apoptosis assay

To visualize apoptotic cells, live zebrafish (7 or 14 dpf) were incubated with acridine orange (5 μg/mL) for 30 min at 28.5°C (in dark), and washed 3x 5 min with embryo water before being embedded in 1% low melting agarose (containing 1:25 tricaine) for confocal live imaging (Leica SP8, 10X air objective, NA=0.4). Average Z-projections were generated, and gray values (intensity) of forebrain, midbrain, and hindbrain were measured using Fiji ImageJ. Data was entered into Prism 9 (GraphPad). Student’s t-test was performed to examine statistical significance.

### Wholemount immunofluorescence

7 dpf zebrafish were genotyped and then fixed with 100% methanol (MeOH) for 20 min at room temperature, followed by storage at -20°C for at least overnight. Samples were gradually brought back to PBSTween (0.1%) using gradient PBST/MeOH wash followed by PBST wash 3x 5 min. Samples were then permeabilized by 5 min H_2_O wash at room temperature, 1 hour 100% acetone incubation at -80°C, and then 5 min H_2_O wash at room temperature. After a brief PBST wash, samples were then blocked (2% goat serum, 1% BSA, 1% DMSO in 1xPBS buffer) for at least 2 hrs at room temperature. Samples were incubated with primary antibodies: anti-acetyl-tubulin (1:200, T7451, Sigma) overnight at 4°C. After 4x 15 min PBST wash, embryos were then incubated with Goat anti-Mouse IgG (H+L) Alexa Fluor 488 (1:150, Cat# A28175, Invitrogen) overnight at 4°C. To examine brain ventricle size, genotyped and PFA fixed 7 dpf zebrafish were directly incubated with DAPI (10 μM in blocking solution) overnight at 4°C. After 4x 15 min PBST wash, embryos were embedded in 1% low melting agarose and imaged using Leica SP8 confocal microscope (40x water immersion, NA=1.10; 1024×1024; step size 0.42 μm). Standard deviation Z-projections were generated and analyzed by Threshold (default) è Binary è Skeletonize Plugin è Analyze Skeleton in Fiji Image J.

### Transmission electron microscopy

Genotyped 14 dpf larvae were anaesthetized using 0.1% tricaine and fixed in Karnovsky’s fixative (2.5% glutaraldehyde/2% PFA in 0.1 M cacodylate buffer, pH 7.5) at room temperature for 2 h, and re-fixed in fresh fixatives overnight at 4°C. The samples were then washed 3×5 min in 0.1 M cacodylate buffer (pH 7.5), post-fixed in 1% osmium (in 0.1 M cacodylate buffer, pH 7.5) for 1.5 h at room temperature, and washed 3×5 min with 0.1 M cacodylate buffer. Samples were then dehydrated with serial ethanol washes (70%, 90%, 95% and 100%), infiltrated with Epon and embedded in Epon to polymerize in a 60°C oven for 24-48 h. Semi-thin (1 µm) and ultra-thin (60 nm) sections were cut using Leica Ultracut ultramicrotomes and transferred on 200 nm formvar-coated copper grids. Grids were post-stained with 2% uranyl acetate at room temperature for 20 min, washed 7×1 min with water and stained with lead citrate for 5 min, followed by 7×1 min water wash. Samples were imaged using a Hitachi HT7800 transmission electron microscope.

## Acknowledgements

The authors wish to thank Xiucheng Cui and Dr. Jason Burgess of SickKids Zebrafish Genetics & Disease Models Core for assistance with generating the *mtmr5* CRISPR/Cas9 deletion line; Ren Li and Dr. Ali Darbandi of SickKids Nanoscale Biomedical Imaging Facility for assistance with TEM sample processing; SickKids Imaging Facility for assistance with confocal microscopy training and maintenance; and SickKids Zebrafish Facility for their diligence in zebrafish care and maintenance.

## Author contributions statement

*Conceptualization* – MZ and JJD. *Methodology* – ML, SG, MZ, and JJD. *Validation* – JL, ML, SG, and MZ. *Formal analysis* – JL, ML, SG, and MZ. *Investigation* – JL, ML, SG and MZ. *Visualization* – JL, ML, SG, and MZ. *Resources* – JJD. *Writing (Original Draft Preparation)* – SG and MZ. *Writing (Review & Editing)* – JL, MZ, and JJD. *Supervision* – SG, MZ, and JJD. *Project administration* – MZ and JJD. *Funding acquisition* – MZ and JJD.

---

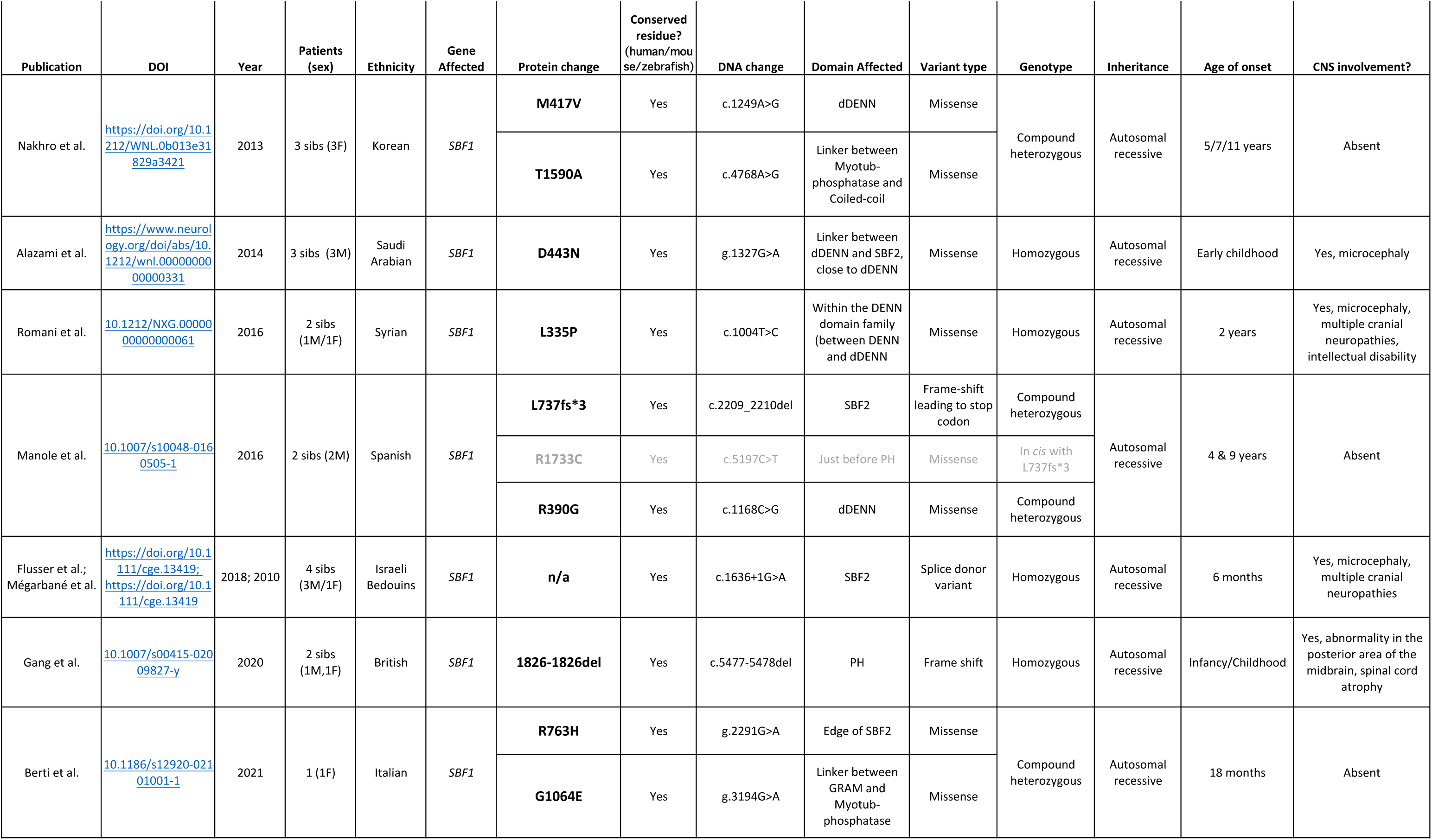
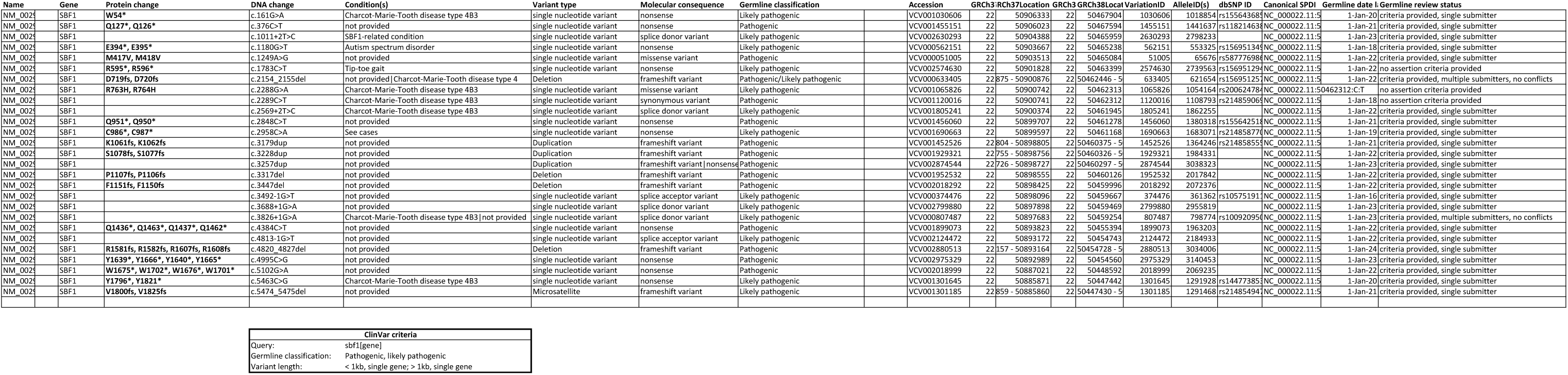
Supp Table 1

